# Wheat *TaNADPO* promotes spot blotch resistance

**DOI:** 10.1101/2024.07.16.602850

**Authors:** Meng Yuan, Qingdong Zeng, Lei Hua, Jianhui Wu, Shuqing Zhao, Mengyu Li, Xiaopeng Ren, Jun Su, Zhuang Ren, Linfei Ma, Zihan Liu, Kaixuan Wang, Manli Sun, Hongfei Yan, Zhensheng Kang, Dejun Han, Shisheng Chen, Xiaodong Wang

## Abstract

*Bipolaris sorokiniana* is a common soil-borne fungal pathogen that can infect various organs of wheat (*Triticum aestivum* L.), causing diseases such as spot blotch, common root rot, head blight, and black point. The genetic basis of wheat resistance to *B. sorokiniana* is not yet fully understood. In this study, a natural population of 1,302 global common wheat germplasms was established and inoculated with *B. sorokiniana* at the seedling stage in a greenhouse. Resistance to spot blotch was assessed, revealing that only about 3.8% of the germplasms exhibited moderate or higher resistance levels. A genome-wide association study (GWAS) using high-density 660K single nucleotide polymorphism (SNP) data identified a region on chromosome 1BL (621.2-674.0 Mb) with 9 SNPs significantly associated (*p* < 10e-4) with spot blotch resistance, designated as *Qsb.hebau-1BL*. RNA sequencing and qRT-PCR assays showed that the gene *TraesCS1B02G410300*, encoding nicotinamide-adenine dinucleotide phosphate-binding oxidoreductase (TaNADPO), was significantly induced by *B. sorokiniana*. Five SNP variations were found in the promoter region of *TaNADPO* in wheat lines with or without *Qsb.hebau-1BL*. Transient expression of *TaNADPO* in *Nicotiana benthamiana* leaves showed a cytoplasmic subcellular localization of the fusion protein with a green fluorescent protein (GFP) tag. Wheat transgenic lines overexpressing *TaNADPO* exhibited significantly enhanced resistance to spot blotch compared to wildtype plants, with higher accumulation of reactive oxygen species (ROS). The knockout EMS mutant of *Triticum turgidum NADPO* (*tdnadpo-K2561*, Gln125*) showed significantly reduced resistance to spot blotch and lower ROS accumulation compared to wildtype plants. In summary, *TaNADPO* has been identified as a crucial gene for resistance to *B. sorokiniana*, providing valuable insights for developing spot blotch-resistant wheat varieties through molecular breeding techniques.

## INTRODUCTION

*Bipolaris sorokiniana*, a fungal pathogen, causes various diseases in wheat and barley, including common root rot, leaf spot blotch, seedling blight, white heads, and grain black point. This pathogen poses a severe threat to wheat production and food security in temperate regions worldwide, with outbreaks more likely under hot and humid conditions (Kumar et al., 2002). The incidence and severity of wheat soil-borne diseases are closely linked to crop rotation systems. Early research indicated that multi-year cereal crop rotations significantly increase the severity of wheat common root rot caused by *B. sorokiniana* (Conner et al., 1996). The large-scale implementation of wheat-maize rotation and straw returning in North China has significantly altered soil organic matter, nutrients, and nitrogen ratios (Wang et al., 2015; Zhao et al., 2006). This rotation system also affects soil microbial diversity and wheat’s disease suppression abilities (Peralta et al., 2018). Consequently, the potential damage from *B. sorokiniana* is increasing annually.

Using disease-resistant varieties remains the most economical and effective method for controlling wheat diseases caused by *B. sorokiniana*. However, current germplasm resources for *B. sorokiniana*-resistant wheat are limited, and research on resistance genetic loci is lagging (Roy et al., 2023; Su et al., 2021). Early studies introduced resistance genes to common root rot from wheat relatives like *Thinopyrum ponticum* (Li et al., 2004). Since resistance is controlled by complex quantitative trait loci (QTL), breeding for *B. sorokiniana* resistance in wheat is challenging (Joshi et al., 2004). Some QTLs have been identified using hybrid populations and linkage mapping, including strong resistance genes *Sb1* on chromosome 7DS (Lillemo et al., 2013), *Sb2* on chromosome 5BL (Kumar et al., 2015), *Sb3* on chromosome 3BS (Lu et al., 2016), *Sb4* on chromosome 4BL (Zhang et al., 2020). Additionally, a minor *Sb* resistance QTL associated with the rust resistance locus *Lr46/Yr29* on chromosome 1BL was detected in the wheat cultivar “Saar” (Lillemo *et al*., 2013).

The completion of the wheat genome sequencing enables the use of the 660K high-density SNP gene chip for genome-wide association studies (GWAS) to explore various quantitative trait genetic loci in wheat (Sun et al., 2020). In GWAS studies on wheat common root rot resistance loci, a US research team used a 15K SNP gene chip and 294 wheat accessions to identify ten pedigrees with strong resistance and six *Sb* resistance QTLs (Ayana et al., 2018). The CIMMYT team used DArTSeq markers for genetic linkage analysis of 301 Afghan wheat varieties resistant to common root rot, finding that about 15% of the Afghan wheat materials had strong resistance, and 25 *Sb* QTL loci were discovered (Bainsla et al.).

This study aims to identify new wheat resistance germplasms and genetic loci associated with spot blotch caused by *B. sorokiniana*. Through omics analysis, we will predict the molecular mechanisms underlying wheat resistance to spot blotch and explore candidate genes controlling this resistance.

## RESULTS

### A spot blotch resistance locus, *Qsb.hebau-1BL*, was identified through phenotyping a large-scale collection of global common wheat germplasms and a genome-wide association study (GWAS)

This collection, consisting of 1,302 released cultivars, breeding lines, and landraces (**Supplemental Figure 1**), was spray-inoculated with *B. sorokiniana* spores at the seedling stage. Resistance to spot blotch was evaluated using a disease severity rating (DSR) scale ranging from 0 to 5 (**Figure 1A** and **Supplemental Table 1**). The results indicated that approximately 74.8% of the wheat germplasms displayed high DSR scores (4-5) to spot blotch (**Figure 1B**), highlighting a scarcity of resistant resources in global common wheat germplasms. Only about 3.8% of the wheat germplasms with DSR scores < 3.0 were classified as resistant (R) or moderately resistant (MR) (**Supplemental Table 2**).

**Figure 1.**
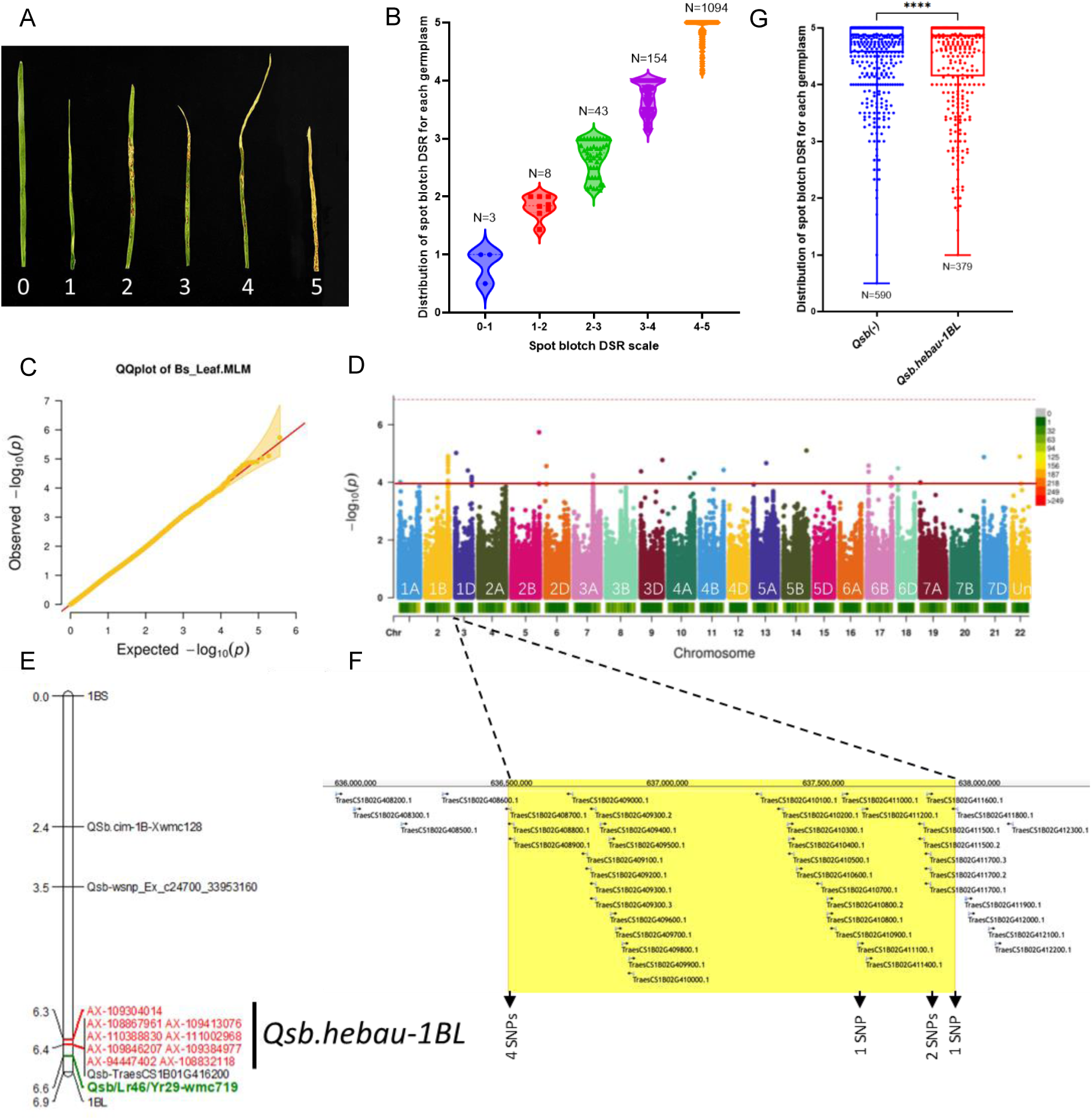
Identification of the genetic locus *Qsb.hebau-1BL* for wheat resistance to spot blotch using GWAS. **(A)** Spot blotch disease severity rating (DSR) in wheat. **(B)** Distribution of spot blotch DSR among the collected global wheat germplasms. **(C)** QQ-plots for GWAS analysis. **(D)** Manhattan plot illustrating associations between SNPs and spot blotch resistance identified through GWAS. **(E)** Distribution map displaying associated SNPs and previously reported spot blotch resistant QTLs on chromosome 1B. **(F)** Physical interval of *Qsb.hebau-1BL* in the Chinese Spring reference genome. **(G)** Phenotypic effects of wheat germplasms with or without *Qsb.hebau-1BL* on spot blotch resistance.

After filtering, a total of 372,277 SNPs were used in the GWAS to detect QTL for spot blotch resistance. The GWAS was conducted using the univariate mixed linear model (MLM) in R software. A total of 34 highly associated SNPs with p < 10e-4 were identified (**Figures 1C** and **1D**, **Supplemental Table 3**). Of these, nine SNPs were clustered in a main genomic region on chromosome 1BL. The SNP-located 621.2-674.0 Mb physical interval on chromosome 1BL was designated as *Qsb.hebau-1BL* (**Figure 1E**). The identified *Qsb.hebau-1BL* was close to a previously reported broad-spectrum resistance QTL to spot blotch, leaf rust, and stripe rust (*Qsb/Lr46/Yr29*). There was a total of 40 highly confident encoded genes within the physical interval of the Chinese Spring reference genome corresponding to *Qsb.hebau-1BL* (**Figure 1F** and **Supplemental Table 4**).

We conducted haplotype analysis on the nine associated SNPs located within the *Qsb.hebau-1BL* interval. By examining the genotypes of resistant and susceptible wheat germplasms, we identified two primary haplotypes, namely *Qsb(-)* and *Qsb.hebau-1BL*, from these nine associated SNPs (**Supplemental Table 5**). Furthermore, we analyzed the genotype of a key SNP, *AX-111002968*, within the *Qsb.hebau-1BL* interval and observed that among the 969 accessions, 379 accessions with the *Qsb.hebau-1BL* haplotype exhibited a significantly lower average DSR value compared to the remaining 590 accessions with the *Qsb(-)* haplotype (**Figure 1G**). This discrepancy in DSR suggests a potential association between the *Qsb.hebau-1BL* haplotype and enhanced resistance to spot blotch.

To validate the findings of GWAS, dCAPS markers associated with *Qsb.hebau-1BL* were designed based on the associated SNPs (**Supplemental Table 6**). Two dCAPS markers were successfully developed to detect the SNP allele in wheat genotypes with or without *Qsb.hebau-1BL* (**Supplemental Figure 2**). These molecular markers provide a reliable tool for identifying the *Qsb.hebau-1BL* locus in different wheat genotypes.

In addition to *Qsb.hebau-1BL*, there were two adjacent SNPs located on chromosome 1DL, two SNPs on chromosome 3AL, two SNPs on chromosome 6BS, and three SNPs on chromosome 6BL, which were also closely linked to the phenotype of wheat resistance to spot blotch. These were temporarily named *Qsb.hebau-1DL* (462.5-462.9 Mb), *Qsb.hebau-3AL* (541.7-542.2 Mb), *Qsb.hebau-6BS* (51.3-51.9 Mb), and *Qsb.hebau-6BL* (671.0-684.8 Mb), respectively (**Supplemental Figure 3** and **Supplemental Table 3**).

### Transcriptome sequencing was initially used to explore the molecular mechanism of wheat resistance to spot blotch

Seedlings of three resistant lines (*R522L_Emai18_Qsb(1BL)*, *R734L_XinXiang9178_Qsb(1BL)*, and *R901_ZhongYu9398_Qsb(-)*) and one susceptible line (*XY22L_Xiaoyan22_Qsb(-)*) were inoculated with either *B. sorokiniana* or water. Twenty-four RNA samples, including three biological replicates for each treatment and genotype combination, were collected at 48 hours post-inoculation (hpi) and subjected to 12-Gb Illumina sequencing (**Supplemental Table 7**). The reference genome of the common wheat “Chinese Spring” v1.1 from the Ensembl Genomes was employed for transcriptome assembly. A total of 143,540 transcripts were assembled and annotated, with transcript expression estimated using FPKM values. In the principal component analysis (PCA) of overall transcript expressions for each sample, most biological replicates clustered together (**Supplemental Figure 4**). All raw sequencing data were deposited at the NCBI under BioProject accession PRJNA1133090.

We initially profiled the expressions of all the pathogenesis-related (*PR*) gene families in the transcriptome (**Figure 2A** and **Supplemental Table 8**). Most *PR* gene families were significantly upregulated upon *B. sorokiniana* infection in both resistant and susceptible lines, indicating an activation of broad-spectrum plant defense responses to the infection of *B. sorokiniana* in wheat. Differentially expressed genes (DEGs) between different combinations of genotype and treatment were identified (|log2-fold change| > 1 and *q*-value < 0.05). A Venn diagram demonstrated the relationships among DEGs (**Figure 2B**). In the susceptible line Xiaoyan22_Qsb(-), there were 8,506 upregulated and 3,274 downregulated DEGs upon *B. sorokiniana* infection. KEGG pathway annotations for DEGs in the comparison of “XY22L_BS vs XY22L_CK” were enriched in metabolisms of glutathione, amino sugar, and phenylalanine (**Figure 2C**). Conversely, in the resistant line *R522L_Emai18_Qsb(1BL)* carrying *Qsb.hebau-1BL*, there were only 2,075 upregulated and 1,121 downregulated DEGs upon *B. sorokiniana* infection. KEGG pathway annotations for DEGs in the comparison of “R522L_BS vs R522L_CK” were enriched in photosynthesis, phenylpropanoid biosynthesis, and mitogen-activated protein kinase (MAPK) signaling pathways (**Figure 2D**). Notably, in the MAPK signaling pathway, clear activations of pattern-triggered immunity (PTI) plant defense response and abscisic acid (ABA) phytohormone were detected (**Supplemental Figure 5** and **Supplemental Table 9**).

**Figure 2.**
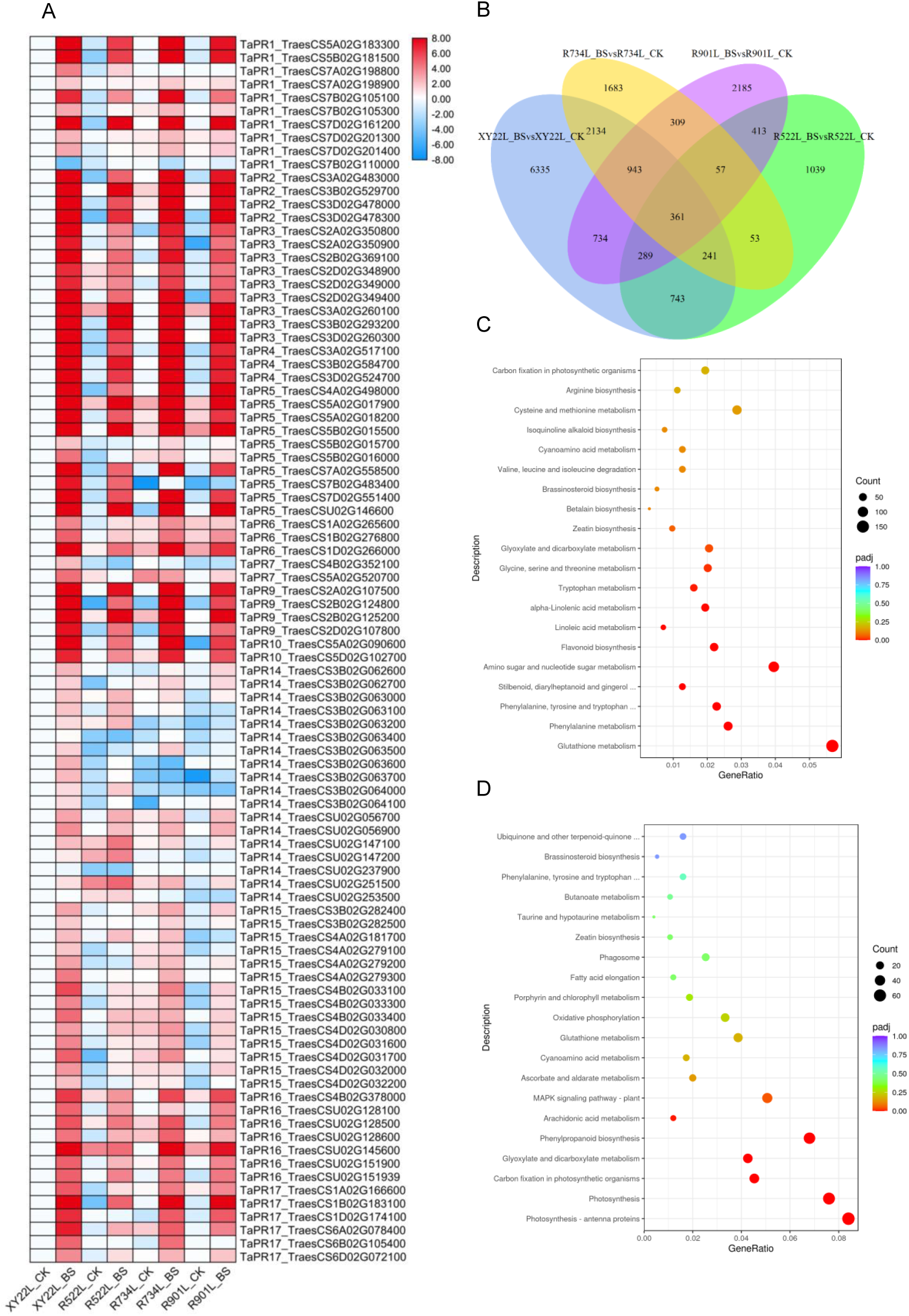
Transcriptome sequencing revealed molecular bases of wheat resistance to spot blotch. **(A)** Expression patterns of *PR* genes in wheat responses to spot blotch. **(B)** Venn diagrams depicting all DEGs between each pair of genotype and treatment combinations. **(C)** Enrichment of KEGG pathways annotated for DEGs in the comparison “XY22_Qsb(-)_BS vs XY22_Qsb(-)_CK”. **(D)** Enrichment of KEGG pathways annotated for DEGs in the comparison “R522_Qsb(+)_BS vs R522_Qsb(+)_CK”.

### *TaNADPO* was identified as the candidate gene within the *Qsb.hebau-1BL* physical interval

The expression profiles of all 40 annotated genes within the *Qsb.hebau-1BL* physical interval were generated based on the RNA-seq database (**Figure 3A** and **Supplemental Table 10**). A gene, *TraesCS1B02G410300*, encoding nicotinamide-adenine dinucleotide phosphate-binding oxidoreductase (TaNADPO), was notably induced by *B. sorokiniana*. To validate the expression pattern of *TaNADPO* in wheat lines with or without *Qsb.hebau-1BL* during *B. sorokiniana* infection, leaf samples from *R_Zhengmai9_Qsb(1BL)* and *S_Xiaoyan22_Qsb(-)* were collected at 24 and 48 hpi with *B. sorokiniana* and subjected to a qRT-PCR assay. The expression level of the *TaNADPO* gene in the wheat line with *Qsb.hebau-1BL* was significantly higher than in the line without *Qsb.hebau-1BL* following *B. sorokiniana* infection at both 24 and 48 hpi (**Figure 3B**).

**Figure 3.**
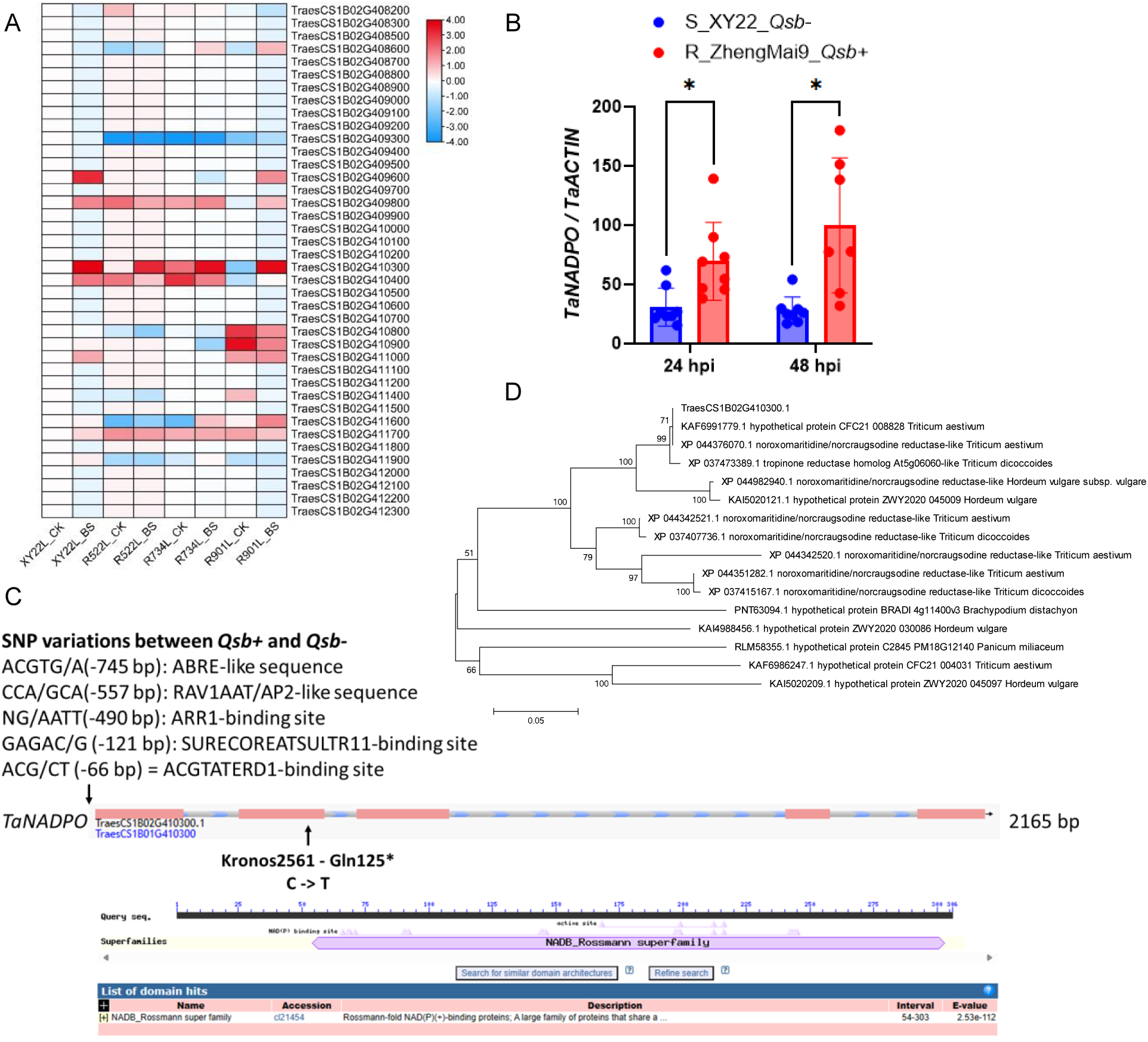
*TaNADPO* was predicted to be the functional gene within the *Qsb.hebau-1BL* locus. **(A)** Expression patterns of all 40 candidate genes within the *Qsb.hebau-1BL* interval during wheat resistance to spot blotch. **(B)** Expression levels of *TaNADPO* in wheat lines with *Qsb.hebau-1BL* (Zheng Mai 9) and without *Qsb.hebau-1BL* (XY22) upon infection with *B. sorokiniana*. **(C)** Gene structure and SNP variations of the *TaNADPO* gene. **(D)** Phylogenetic tree of TaNADPO protein and its homologs in other plant species.

To explore the variation of the *TaNADPO* gene among different lines, the genomic region of the *TaNADPO* gene was amplified from 10 resistant lines with *Qsb.hebau-1BL* and 10 susceptible lines without *Qsb.hebau-1BL* (**Supplemental Table 5**). Five SNP variations were identified at positions −745, −557, −490, −121, and −66 bp in the promoter region of the *TaNADPO* gene in wheat lines with or without *Qsb.hebau-1BL*. These variations are located within the binding sites of ABRE-like, RAV1AAT/AP2-like, ARR1, SURECOREATSULTR11, and ACGTATERD1, respectively (**Figure 3C** and **Supplemental Table 11**). Three gene-specific dCAPS markers were developed based on these SNPs in the promoter region of the *TaNADPO* gene (**Supplemental Figure S6** and **Supplemental Table 12**). We were unable to locate any homologous copies of *TaNADPO* within the A and D sub-genomes of common wheat. The encoded TaNADPO protein includes a predicted domain of the NADB_Rossmann superfamily (**Figure 3C**) and is conserved across various plant species (**Figure 3D**).

### The transient expression of the *TaNADPO* gene in tobacco leaves significantly enhanced plant resistance to *B. sorokiniana* and the generation of ROS

To determine the subcellular localization of TaNADPO, a green fluorescent protein (GFP) tag was utilized to visualize TaNADPO in *Nicotiana benthamiana* leaves. Cytoplasmic fluorescence was observed in tobacco leaves for both GFP-TaNADPO and TaNADPO-GFP constructs (**Figure 4A**). Nitroblue tetrazolium (NBT) was used to visualize the accumulation of superoxide anions (O^2-^) in the *Agrobacterium*-infiltrated *N. benthamiana* leaves (**Figure 4B**). The percentage of the stained area in each fully infiltrated leaf was calculated using ASSESS software. The accumulation of O^2-^ in the *TaNADPO*-expressed *N. benthamiana* leaves was significantly higher than in the GFP control (**Figure 4C**).

**Figure 4.**
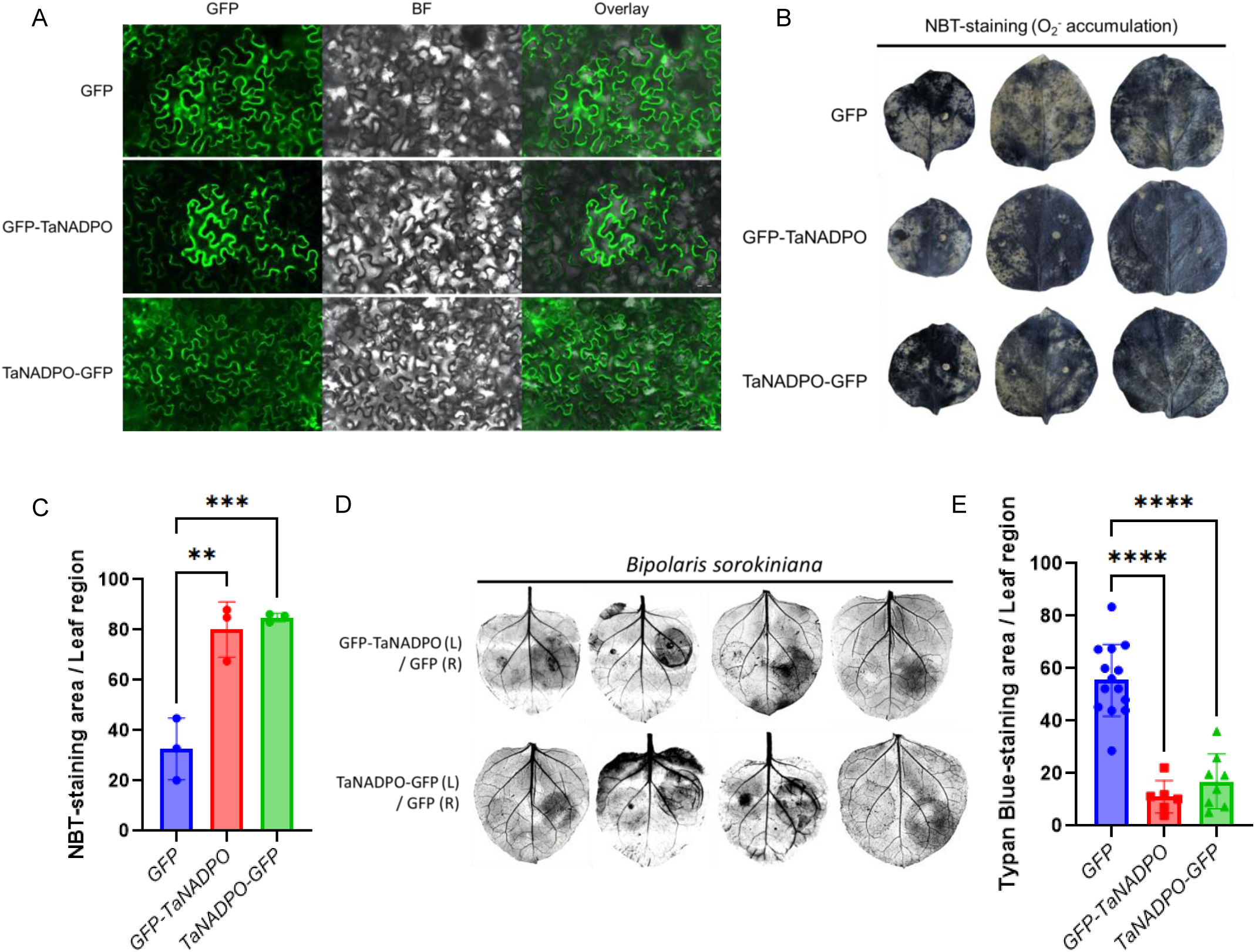
Initial functional characterization of GFP-labeled TaNADPO in *Nicotiana benthamiana*. **(A)** Subcellular localization of GFP-TaNADPO and TaNADPO-GFP transiently expressed in leaves of *Nicotiana benthamiana*. Scale bar = 50 μm. **(B)** Accumulation of ROS triggered by infection of *Agrobacterium tumefaciens* in leaves of *N. benthamiana* expressing GFP-TaNADPO and TaNADPO-GFP. Empty GFP was used as a control. **(C)** Proportions of NBT-staining areas in the whole leaf region estimated using ASSESS software and employed for statistical analysis. **(D)** Infection of *Bipolaris sorokiniana* in the leaf region transiently expressing GFP-TaNADPO and TaNADPO-GFP. Empty GFP was used as a control. **(E)** Proportions of trypan blue-stained cell death areas in the leaf region pre-infiltrated with *A. tumefaciens* carrying *GFP-TaNADPO* and *TaNADPO-GFP*, estimated using ASSESS software and employed for statistical analysis.

To initially investigate the roles of *TaNADPO* in plant resistance to *B. sorokiniana*, *TaNADPO* was transiently expressed in *N. benthamiana* leaves, followed by inoculation with *B. sorokiniana* at 2 days post-infiltration. Plant cell death was evaluated by measuring the proportion of the Typan Blue staining area in the infiltrated leaf region (**Figure 4D**). Compared to leaves infiltrated with free *GFP*, the expression of either *GFP-TaNADPO* or *TaNADPO-GFP* significantly enhanced plant resistance to *B. sorokiniana* (**Figure 4E**).

### The transgenic wheat line overexpressing the *TaNADPO* gene exhibited increased resistance to spot blotch due to a heightened burst of ROS

Transgenic wheat lines overexpressing *TaNADPO* (*TaNADPO-OE*) were cultivated, and the *TaNADPO* transgene displayed high expression levels across different lines (**Figure 5A**). The wheat transgenic lines *TaNADPO-OE* demonstrated markedly enhanced resistance to spot blotch in comparison to wildtype plants (**Figure 5B**). Upon inoculation with *B. sorokiniana*, the leaves of wildtype plants exhibited full necrosis at 10 days post-inoculation (dpi), while those of *TaNADPO-OE* only displayed small lesions. The percentage of the spot blotch region in each inoculated leaf was analyzed, revealing a significant reduction in *B. sorokiniana* infection in *TaNADPO-OE* (**Figure 5C**).

**Figure 5.**
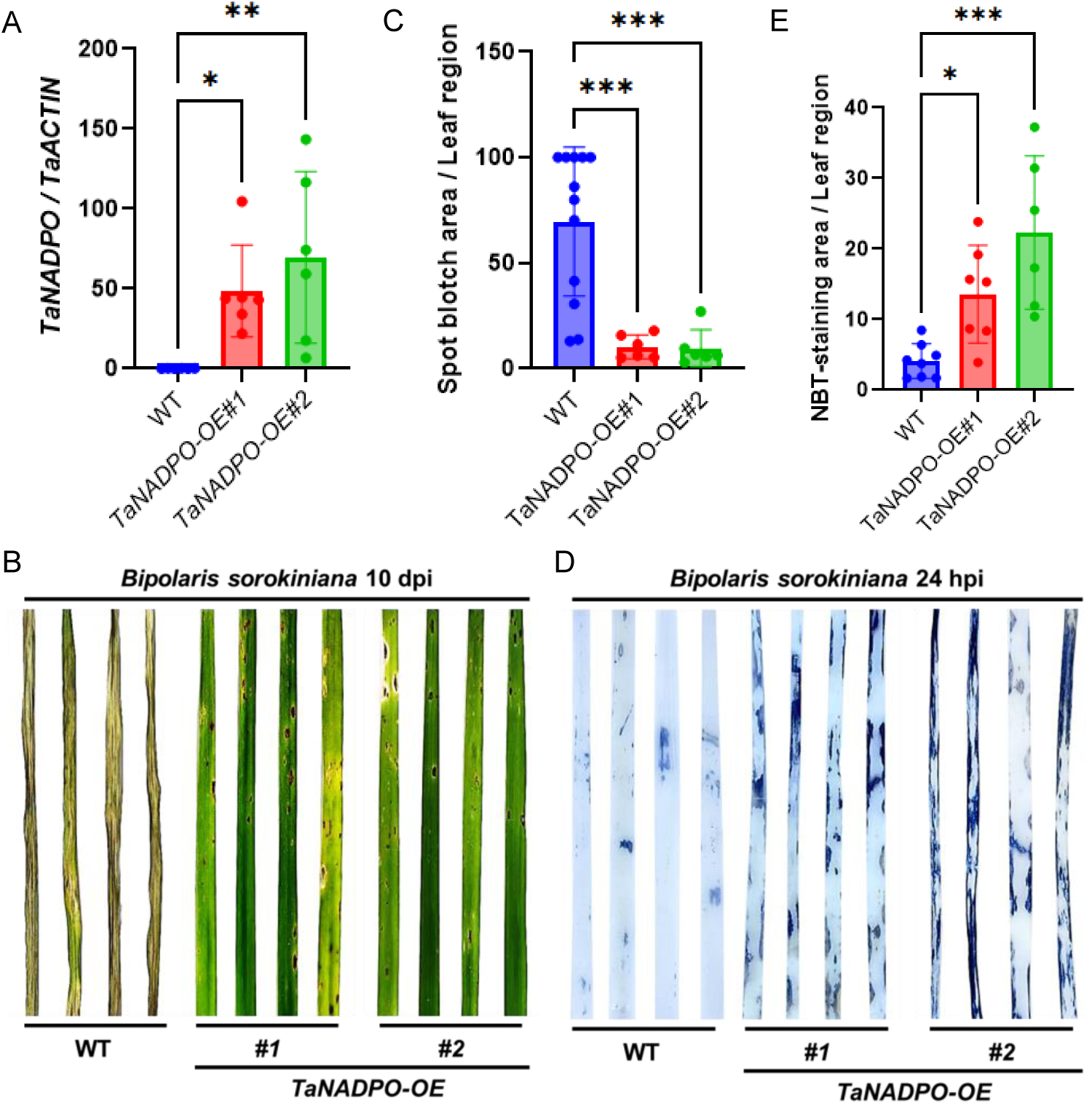
*TaNADPO* promotes wheat resistance to spot blotch with higher accumulation of ROS. **(A)** Transcript levels of the *TaNADPO* transgene in *TaNADPO-OE*, determined by RT-qPCR. **(B)** Phenotypes of *TaNADPO-OE* and wild-type plants in response to *B. sorokiniana*. **(C)** Statistical analysis of the proportion of the spot blotch region on leaves of *TaNADPO-OE* and wild-type plants. **(D)** NBT-staining of leaves from *TaNADPO-OE* and wild-type plants inoculated with *B. sorokiniana* at 24 hours post-inoculation. **(E)** Statistical analysis of the proportion of the NBT-staining region on leaves of *TaNADPO-OE* and wild-type plants infected with *B. sorokiniana*.

Furthermore, the accumulation of O^2-^ induced by *B. sorokiniana* infection at 24 hours post-inoculation was visualized using NBT staining (**Figure 5D**). Leaves of *TaNADPO-OE* accumulated a significantly higher amount of O^2-^ upon *B. sorokiniana* infection compared to wildtype plants (**Figure 5E**).

### The knock-out line *tdnadpo-K2561* of tetraploid wheat showed increased susceptibility to spot blotch, along with reduced accumulation of ROS

In the tetraploid wheat (*Triticum turgidum* ssp. *durum*), the TaNADPO homolog, TdNADPO, exhibited a protein similarity of 99.3% with TaNADPO (as shown in the sequence alignment in **Supplemental Figure S7**). The mutant K2561, a stop-gained EMS mutant of *TdNADPO* in the background of tetraploid wheat Kronos, was obtained from a previous exome capture project on durum wheat. The predicted point mutation of *TdNADPO* at nucleotide position 373, resulting in a CAG to TAG in-frame mutation, led to the gain of a stop codon, replacing the encoded Gln at position 125. This mutation in the K2561 mutant line was confirmed through PCR amplification and Sanger sequencing (**Figure 6A**).

**Figure 6.**
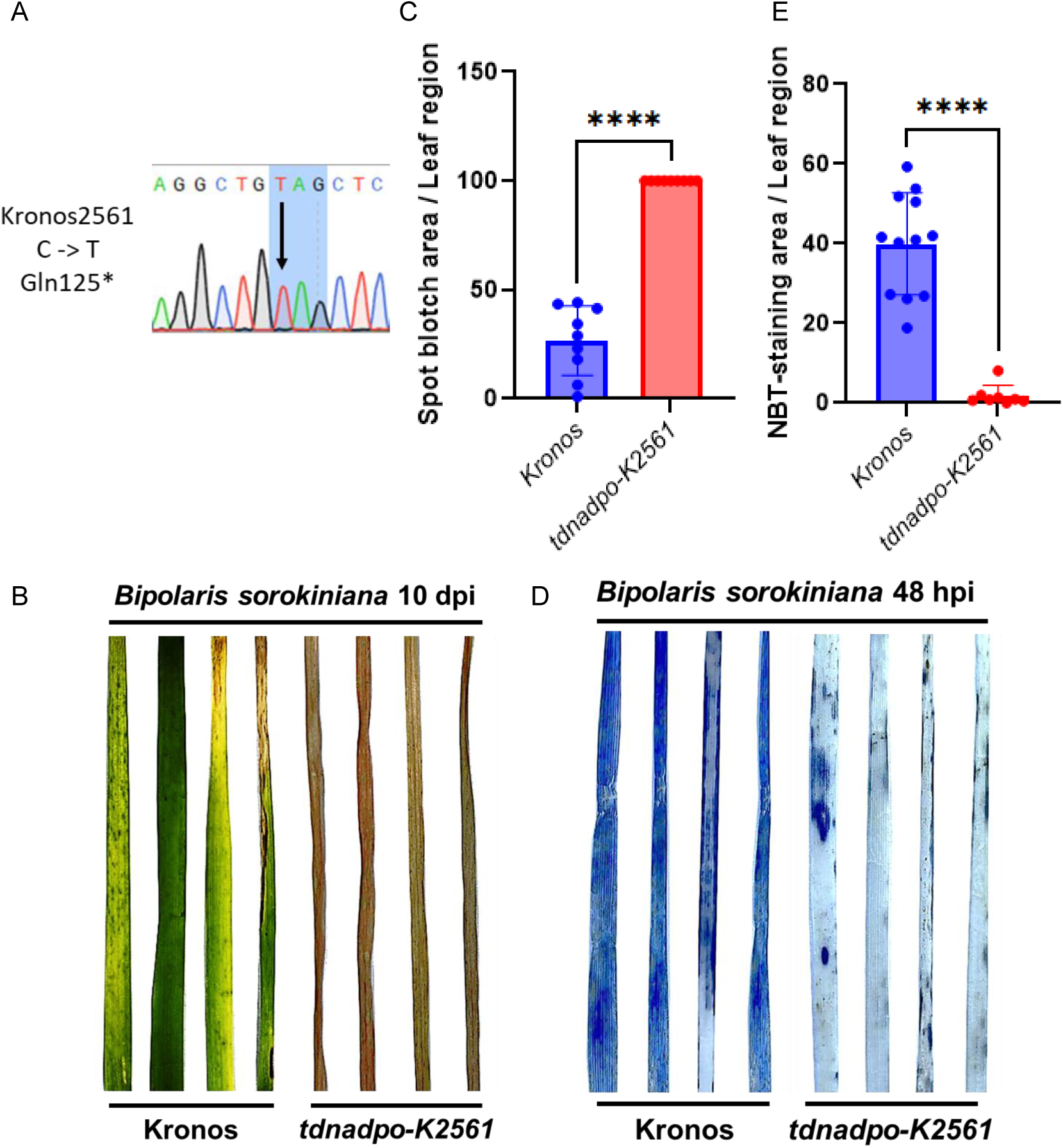
Tetraploid wheat *tdnadpo* knock-out mutant showed diminished resistance to spot blotch and attenuated ROS generation. **(A)** Sanger sequencing validation of the stop-gained mutation in *TdNADPO* in the Kronos EMS mutant K2561. **(B)** Phenotypes of *tdnadpo-K2561* and wild-type Kronos plants in response to *B. sorokiniana*. **(C)** Statistical analysis of the proportion of the spot blotch region on leaves of *tdnadpo-K2561* and wild-type plants. **(D)** NBT-staining of leaves from *tdnadpo-K2561* and wild-type Kronos plants inoculated with *B. sorokiniana* at 48 hpi. **(E)** Statistical analysis of the proportion of the NBT-staining region on leaves of *tdnadpo-K2561* and wild-type plants infected with *B. sorokiniana*.

Subsequently, the knockout line *tdnadpo-K2561* and its wildtype Kronos were subjected to spray-inoculation with *B. sorokiniana* (**Figure 6B**). While the wildtype Kronos exhibited relatively higher resistance to spot blotch compared to common wheat materials, the *tdnadpo-K2561* knockout line showed significantly reduced resistance to spot blotch when compared to the wildtype plants. The lesion region percentage in the inoculated leaves of *tdnadpo-K2561* was notably higher than that in the wildtype plants (**Figure 6C**). Furthermore, visualization of O^2-^ accumulation using NBT staining at 48 hours post-inoculation (hpi) revealed that the knockout line *tdnadpo-K2561* exhibited significantly lower levels of ROS production compared to the wildtype plants upon *B. sorokiniana* infection (**Figures 6D** and **6E**).

## DISCUSSION

*Bipolaris sorokiniana* can infect various organs of the wheat plant throughout its growth stages, leading to spot blotch in leaves, common root rot in stems and roots, white heads in spikes, and black points in grains (Al-Sadi, 2021). It is noteworthy that *Bipolaris oryzae*, a relative in the same genus causing brown spot disease in rice, poses a significant threat to rice production in India and Bangladesh (Barnwal et al., 2013). Given these concerns, especially in the context of climate change, developing resistance to *B. sorokiniana* is crucial in wheat breeding programs, particularly in warm and humid regions.

Gupta *et al*. summarized the primary sources of resistance against spot blotch in global wheat germplasm (Gupta et al., 2018). Resistance sources against spot blotch appear to be limited in the global wheat germplasm pool. Only approximately 3.8% of collected global wheat germplasms with DSR scores < 3.0 were identified as potential resistance sources in this study. Similarly, a previous investigation into 294 genotypes of hard winter wheat found that only about 5.2% exhibited high resistance (Ayana *et al*., 2018). These findings underscore the urgent need to explore novel genetic loci that control resistance to spot blotch and integrate newly identified resistance sources into breeding practices.

According to our previous review, as of 2021, a total of 85 genetic loci located across 19 chromosomes of wheat have been identified to regulate plant resistance to *B. sorokiniana* infection (Su *et al*., 2021). Among these loci, *Sb1* on chromosome 7DS is linked to the broad-spectrum resistance gene *Lr34/Yr18/Pm38*, which encodes an ATP-binding cassette (ABC) transporter that imparts resistance to leaf rust, stripe rust, and powdery mildew (Krattinger et al., 2009). Other dominant *Sb* genes, namely *Sb2*, *Sb3*, and *Sb4*, have not yet been cloned. In the present study, employing a GWAS approach, 9 closely associated SNPs were found to be significantly linked to resistance against spot blotch on chromosome 1BL. The identified genetic locus, designated *Qsb.hebau-1BL*, is in close proximity to a previously recognized minor *Sb* resistance QTL associated with the rust resistance locus *Lr46/Yr29* (Lillemo *et al*., 2013). The *Lr46/Yr29*, which governs slow rusting resistance, has not been cloned to date. Exploring candidate genes within *Qsb.hebau-1BL* may yield valuable insights for the eventual discovery of *Lr46/Yr29* in the future.

While only a limited number of *Sb* genes have been cloned, several transcriptome studies have initially delved into the molecular mechanisms underlying wheat resistance to *B. sorokiniana* infection, encompassing plant responses to spot blotch (Zhang et al., 2022), common root rot (Qalavand et al., 2023), and black point (Li et al., 2021). Our transcriptome analysis revealed that susceptible responses to spot blotch were linked to nutrient metabolism, whereas resistant responses to spot blotch were associated with the activation of *PR* genes, phenylpropanoid biosynthesis, and MAPK signaling in the PTI pathway. Similar to other typical necrotrophic pathogens, *B. sorokiniana* likely secretes polygalacturonases (PGs) during early stages of infection, targeting the homogalacturonan component of pectin and releasing oligogalacturonides (OGs) of varying chain lengths. These OGs are known to trigger typical PTI responses, such as oxidative burst, accumulation of phytoalexins, and hormone biosynthesis (Ghozlan et al., 2020).

Based on the gene expression and SNP variations of the candidate *TaNADPO* gene among different genotypes with or without *Qsb.hebau-1BL*, the functions of *TaNADPO* in plant resistance to *B. sorokiniana* infection and the accumulation of ROS were characterized in both *N. benthamiana* and wheat. Nicotinamide adenine dinucleotide (NAD) and its phosphorylated variant, nicotinamide adenine dinucleotide phosphate (NADP), play pivotal roles as energy transducers and signaling molecules across all forms of life. Within plants, these pyridine nucleotides are integral to various processes, encompassing core energy metabolism, growth, and immune responses (Smith et al., 2021). Furthermore, NAD also serves as a key component in plant defense-related signaling and responses, including ROS generation, calcium signaling, DNA repair, and protein deacetylation (Pétriacq et al., 2013). In the wheat genome, a total of 46 *NADPO* family genes have been identified, playing widespread roles in plant responses to a broad range of environmental stresses (Hu et al., 2018). However, only a few *TaNADPO* genes have been functionally validated during wheat resistance to phytopathogens.

In the current study, we found that spot blotch resistance controlled by the *TaNADPO* gene was associated with its expression levels in different genotypes. Five SNPs identified in the promoter region of the *TaNADPO* gene might influence the binding efficiency of corresponding transcription factors. Future research will focus on exploring the upstream regulatory network of the *TaNADPO* gene. Nevertheless, the developed dCAPS markers for either the *Qsb.hebau-1BL* genetic locus or the *TaNADPO* gene will greatly facilitate molecular breeding for wheat resistance to spot blotch.

In conclusion, in a study involving inoculation with *B. sorokiniana*, only 3.8% of the 1,302 tested wheat lines showed moderate or higher resistance to spot blotch. A GWAS identified the *Qsb.hebau-1BL* region on chromosome 1BL with significant SNP associations for resistance. The *TaNADPO* gene, encoding nicotinamide-adenine dinucleotide phosphate-binding oxidoreductase, was significantly induced by *B. sorokiniana*. Wheat lines overexpressing *TaNADPO* showed increased resistance and higher ROS accumulation. Conversely, the EMS mutant *Triticum turgidum tdnadpo* had reduced resistance and lower ROS levels. *TaNADPO* is crucial for resistance to *B. sorokiniana*, providing valuable insights for breeding spot blotch-resistant wheat.

## METHODS

### Plant materials and spot blotch inoculation

**(A)** *B. sorokiniana* was cultured on potato dextrose agar (PDA) medium at 28 °C. Fungal spores were collected from the cultured dishes by gently scraping the mycelium into a sterile water solution. The collected solution was then filtered using sterile gauze to obtain a spore suspension for inoculation. The spore suspension concentration was adjusted to 10^5^/mL using a microscope and hemocytometer.

A mixture of nutrient soil and vermiculite in a 2:1 ratio was used for growing the wheat seedlings. Eight full-grain seeds of each wheat line were planted in 32-hole trays. The wheat seedlings were grown in a greenhouse at 20-22 °C with a photoperiod of 16 hours light and 8 hours dark. At the two-leaf stage, the seedlings were inoculated by spraying *B. sorokiniana* spores and then transferred to a dark dew chamber with 100% humidity at 20-22 °C for 48 hours. Approximately 10 days post-inoculation (dpi), disease symptoms of spot blotch were recorded for each plant using a disease severity rating (DSR) scale ranging from 0 to 5:

- Level 0: No disease spots on wheat leaves.
- Level 1: Sporadic disease spots with small areas on wheat leaves.
- Level 2: Disease spots with larger areas on wheat leaves.
- Level 3: Disease spots covering about 30% of the total leaf area, with necrosis at the leaf tips.
- Level 4: Disease spots covering about 50% of the total leaf area, with necrosis at the leaf tips.
- Level 5: Disease spots covering more than 70% of the total leaf area, or complete withering of the entire leaf.

The DSR for each wheat line was calculated based on the severity data of individual plants. Wheat lines underwent 2-3 independent repetitions of *B. sorokiniana* inoculation, and the final data were obtained by integrating these repetitions to determine the spot blotch resistance phenotype of the entire population.

### Genotyping and association analysis

The 1,302 common wheat germplasm resources were genotyped using a 660K high-density SNP chip as described (Wu et al., 2021). Briefly, after obtaining the genotype data, the Affymetrix Genotyping Console™ (GTC) software was used for initial data processing and analysis. SNP loci with a genotype missing rate greater than 10% were deleted; loci with a minor allele frequency less than 5% were removed; and loci failing the Hardy-Weinberg equilibrium test at a 1% threshold were excluded. The processed SNP data were then used to perform a genome-wide association study (GWAS). Additionally, the Polymorphic Information Content (PIC) was calculated to reflect the diversity of the variable SNP loci.

Population structure analysis of the 1,302 wheat materials was conducted using Structure v2.2.3 software based on the Bayesian algorithm. The burn-in parameter was set to 10,000, and the Monte Carlo Markov Chain (MCMC) was set to 20,000, with the analysis repeated 5 times. The range of subgroup number (K) was from 2 to 10, and the optimal (K) value was determined based on ΔK. The kinship relationships were represented by a kinship heatmap drawn using R (**Supplemental Figure 8**).

Linkage disequilibrium (LD) analysis of the whole genome and sub-genomes was conducted using PLINK software. LD was calculated with R^2^ as the reference standard for pairwise markers after SNP data processing. The step-shift window was set to 1,000 Kb, and the sliding window was set to 30,000 Kb. When drawing the distance decay LOESS curve, the genome LD decay distance standard was the marker interval when R^2^ ≤ 0.1. The LD decay graph of the wheat genome included the LD decay distances of the whole wheat genome and the ABD sub-genomes (**Supplemental Figure 9**). High-confidence loci within the same chromosome LD decay distance were considered the same QTL locus.

Using a univariate mixed linear model, R software was employed to perform association analysis of the spot blotch phenotypic data of the 1,302 wheat lines with 660K high-density SNP genotype data. GEC software calculated the *P*-value range for each chromosome, and 10^-4^ was regarded as the threshold for significant loci based on the association analysis results. The results were visualized using R’s QQman package to draw Manhattan plots and QQ plots. Additionally, GCTA software was used to calculate the genetic contribution of loci on chromosomes.

### Genetic loci analysis and dCAPS marker design

The chromosome positions of the significantly associated SNPs and nearby genes were analyzed using the common wheat Chinese Spring reference genome (v1.1). The genetic loci identified in this study were mapped to chromosomes using MapChart v2.32 software and compared with previously reported loci on the same chromosome (Su *et al*., 2021). Haplotype analysis was performed for the clustered significant SNPs, followed by phenotypic effect analysis based on the haplotype and phenotype data of the GWAS population.

Based on the GWAS results, SNPs closely linked to *Qsb.hebau-1BL* were selected to develop derived cleaved amplified polymorphic sequence (dCAPS) markers (**Supplemental Table 6**) following the method described (Neff et al., 1998). The NEBcutter tool v2.0 [http://www.labtools.us/nebcutter-v2-0/] was used to identify potential restriction sites at each SNP. Genome-specific primers were designed using the Primer 3 website [https://bioinfo.ut.ee/primer3-0.4.0/primer3/] to amplify regions flanking the target SNPs. PCR products were sequenced to validate the presence of the SNPs. The amplified PCR products were then digested using specific restriction enzymes. The reliability of the molecular markers was assessed by analyzing the correlation between the detected genotypic signals and disease resistance phenotypes.

### RNA-sequencing and bioinformatic analysis

Seedlings of selected wheat lines resistant to *B. sorokiniana*, namely R522 (EMai 18), R734 (XinXiang 9178), and R901 (ZhongYu 9398), along with susceptible control XY22 (XiaoYan 22), were spray-inoculated with *B. sorokiniana* spores as described previously. Sterilized water was used as a mock inoculation control (CK). At 5 dpi, leaf samples were collected and sent to Novogene Corporation Ltd. for RNA sequencing. For each genotype-treatment combination, three biological replicates were prepared, and RNA samples were sequenced using the Illumina HiSeq 1000 System with a target of 12 Gb per sample. Transcriptome assembly was performed using Hisat2 v2.2.1 software (Kim et al., 2019) with the reference genome of the common wheat cultivar “Chinese Spring” v1.1. Gene expression levels were estimated using the expected number of fragments per kilobase of transcript sequence per million base pairs (FPKM) with StringTie v1.3.3b software (Pertea et al., 2015). Expression of PR genes was determined based on gene accessions provided in a previous study (Zhao et al., 2024). Heatmaps based on FPKM values were created using TBtools v1.108 software (Chen et al., 2020). Differentially-expressed genes (DEGs) were identified using DESeq2 v1.22.1 software (Love et al., 2014), and Gene Ontology (GO) (Young et al., 2010) and Kyoto Encyclopedia of Genes and Genomes (KEGG) (Kanehisa et al., 2007) annotations were applied for DEG annotation.

### Expression and variations of the *TaNADPO* gene

Seedlings of the wheat resistance line ZhengMai 9, which carries *Qsb.hebau-1BL*, and the susceptible line XiaoYan 22, which lacks *Qsb.hebau-1BL*, were spray-inoculated with *B. sorokiniana* spores as described previously. Samples were collected from inoculated leaves at 24 and 48 hours post-inoculation (hpi). RNA was extracted using the RNA Extraction Kit (QIAGEN, Hilden, Germany), and first-strand cDNA was synthesized using the Reverse Transcription Kit (Takara, Dalian, China). Primers for the qRT-PCR assay targeting the *TaNADPO* gene were designed (**Supplemental Table 13**). The wheat reference gene *TaActin* (GenBank accession AB181991) was used as an endogenous control (Paolacci et al., 2009). Transcript levels were expressed relative to *TaActin* using the 2^−ΔCt^ method as described (Li et al., 2023). Data analysis and multiple unpaired t-tests were conducted using Prism software v10.0.

The genomic region of the *TaNADPO* gene was amplified from 10 resistance lines carrying *Qsb.hebau-1BL* and 10 susceptible lines lacking *Qsb.hebau-1BL* (**Supplemental Table 14**). The amplified DNA was sequenced using Sanger sequencing, and variations in the promoter region of the *TaNADPO* gene were identified. Gene-specific dCAPS markers for the *TaNADPO* gene were designed based on these SNPs (**Supplemental Table 12**).

Homologous sequences of the wheat *TaNADPO* protein were obtained from wheat and its wild relatives’ reference genomes [https://wheat.pw.usda.gov/blast/] and the NCBI database [https://www.ncbi.nlm.nih.gov]. Multiple sequence alignment was performed using the MUSCLE method in MEGA software v7.0. The phylogenetic tree was generated using the pair-wise deletion method with bootstrap values based on 1,000 iterations and visualized using Interactive Tree of Life (iTOL) v5.0 [https://itol.embl.de/]. The conserved domain of the deduced *TaNADPO* protein was predicted using the NCBI Structure website [https://www.ncbi.nlm.nih.gov/Structure/cdd/wrpsb.cgi].

### Transient expression assay in tobacco leaves

The coding regions of the full-length TaNADPO protein were inserted into both pGWB5 (*35S::gene-GFP*) and pMDC43 (*35S::GFP-gene*) vectors. These recombinant constructs were expressed in *Nicotiana benthamiana* leaves using transformed *Agrobacterium tumefaciens* strain GV3101. Fluorescence was examined 48 hours post-infiltration using a Nikon Ti-2 microscope.

For reactive oxygen species (ROS) staining, tobacco leaves infiltrated with transformed *A. tumefaciens* were collected at 1 dpi and stained using nitroblue tetrazolium (NBT) to visualize the accumulation of superoxide anion (O^2−^) as described (Zhao et al., 2020). Briefly, tobacco leaves were collected and soaked in a solution containing 10 mM NaN_3_ and 10 mM potassium phosphate buffer (pH 7.8) with 0.1% NBT (w/v), or in a hydrochloric acid (HCl)-acidified solution (pH 3.8) containing 1 mg/mL of DAB for 24 hours. Samples were then decolorized in boiling 95% ethanol for 10 minutes. The percentage of stained area in each leaf was determined using

ASSESS 2.0 software (Lamari, 2008), and statistical analysis was conducted using one-way ANOVA with Prism software v10.0. For the *B. sorokiniana* inoculation assay, two days post-infiltration of transformed *A. tumefaciens*, the inoculation solution of *B. sorokiniana* spores was infiltrated into the same area as described (Wang et al., 2024). Approximately three days later, disease lesions/necrosis induced by *B. sorokiniana* on tobacco leaves were observed and evaluated using ASSESS 2.0 software.

### Wheat transgenic line overexpressing *TaNADPO*

Transgenic wheat lines used in this study were generated in our lab with technical assistance from Jinan Bangdi Ltd. Company, Shandong, China. The transgenic background material “JW1” was selected by the company from a segregating population resulting from the cross between spring common wheat cultivars “Fielder” and “NB1”. To create the transgenic wheat line *TaNADPO-OE*, the full-length ORF of *TaNADPO* was cloned into the wheat transgenic vector pLGY-02, which contains the maize (*Zea mays*) ubiquitin 1 promoter and T-DNA insertion site. *Agrobacterium*-mediated transformation was employed to generate the transgenic lines. Initial validation of transgenic lines was performed by PCR amplification of the transgene insertion in genomic DNA using specific primers (**Supplementary Table S13**). The expression levels of the *TaNADPO* transgene in the wheat transgenic lines were determined by qRT-PCR assay. Statistical analysis was conducted using one-way ANOVA with Prism software v10.0.

For evaluation of disease resistance, seedlings of the transgenic wheat line *TaNADPO-OE* and wild-type plants were maintained and inoculated with *B. sorokiniana* following established procedures and conditions. The fully expanded third leaves of wheat seedlings were spray-inoculated with *B. sorokiniana* spores in a water solution. Disease symptoms were assessed at 10 dpi, and the percentages of necrotic areas on each inoculated leaf were quantified using ASSESS image analysis software v2.0 for plant disease quantification (Lamari, 2008).

To visualize the accumulation of reactive oxygen species (ROS) in *B. sorokiniana*-infected wheat leaves, NBT was used to stain O^2−^ in wheat leaves (Zhao et al., 2021). Leaf samples spray-inoculated with *B. sorokiniana* were collected at 24 hpi. For O^2−^ visualization, samples were soaked in a solution of 10 mM sodium azide (NaN_3_) and 10 mM potassium phosphate buffer (pH 7.8) containing 0.1% NBT (w/v) for 24 hours. Samples were then decolorized in boiling 95% ethanol for 10 minutes. The percentage of stained area in each leaf was determined using ASSESS software v2.0 (Lamari, 2008).

The experiment included two independent transgenic lines, and each line consisted of 6-10 biological replicates. Statistical analysis was performed using Prism software v10.0, and one-way ANOVA was conducted to determine significant differences.

### EMS mutant of tetraploid wheat *tdnadpo* knock-out line

The homolog of *TaNADPO* in tetraploid wheat (*TdNADPO*) was identified using annotations in the JBrowse database for wheat genomes available on the WheatOmics website [http://wheatomics.sdau.edu.cn/jbrowse.html]. The protein sequences of TaNADPO and TdNADPO were aligned using MEGA software v7.0. A stop-gained EMS mutant of the *TdNADPO* gene was predicted from a previous exome capture project on Kronos (Krasileva et al., 2017). This mutant line, K2561, was obtained from the publicly released seed library at Shandong Agricultural University, China. The genomic region containing the predicted mutation at 373 nt from the initiation codon of the *TdNADPO* gene was amplified from both K2561 and wild-type Kronos plants. The amplified DNA was sequenced using Sanger sequencing.

Seedlings of the *tdnadpo*-K2561 knockout line and wild-type plants were spray-inoculated with *B. sorokiniana*. Accumulation of O^2−^ was visualized using NBT staining at 48 hpi, and disease symptoms were assessed at 10 dpi. ASSESS image analysis software v2.0 was used to analyze the percentage of spot blotch area or NBT-stained area in the leaf region. Each line consisted of 6-10 biological replicates. Statistical analysis was performed using Prism software v10.0, and an unpaired t-test was conducted to determine significant differences.

## CRediT authorship contribution statement

X.W., S.C., and D.H. designed the research; M.Y., Q.Z., L.H., J.W., S.Z., M.L., X.R., J.S., Z.R., L.M., Z.L., and K.W. performed the research; M.S., H.Y., and Z.K. analyzed the data; X.W. wrote the manuscript. All authors reviewed the manuscript.

## Acknowledgments

We would like to thank Prof. Jiajie Wu from Shandong Agricultural University for sharing the Kronos EMS mutants. Work at XW laboratory was supported by the National Key Research and Development Program of China (2023YFD1201002), Local Science and Technology Development Fund Projects of Hebei Province Guided by the Central Government (236Z6501G), Provincial Natural Science Foundation of Hebei (C2022204010), Key Research and Development Project of Shijiazhuang City for University in Hebei Province (241490012A), Hebei Province Dryland Alkali-resistant Wheat Industry Technology System (HBCT2024030206), State Key Laboratory of North China Crop Improvement and Regulation, and S&T Program of Hebei (23567601H). Work at SC laboratory was supported by the Key R&D Program of Shandong Province (ZR202211070163).

## Appendix A. Supplementary data

**Supplemental Figure 1.** Types of wheat germplasms in the studied natural population.

**Supplemental Figure 2.** Validation of dCAPS markers designed based on SNPs within the *Qsb.hebau-1BL* locus.

**Supplemental Figure 3.** Distribution map displaying associated SNPs and previously reported spot blotch resistant QTLs on chromosomes other than Chr1B.

**Supplemental Figure 4.** Principal component analysis (PCA) of the overall transcript expressions for each sample in the transcriptome.

**Supplemental Figure 5.** DEGs in comparison of “R522L_BS vs R522L_CK” enriched in the KEGG pathway of “MAPK signaling pathway – plant”.

**Supplemental Figure 6.** Development and validation of three dCAPS markers based on variations in the *TaNADPO* gene.

**Supplemental Figure 7.** Protein sequence alignment of TaNADPO with TdNADPO.

**Supplemental Figure 8.** Population structure analysis of the studied natural population.

**Supplemental Figure 9.** Linkage disequilibrium (LD) decay map of the SNPs utilized in this study.

**Supplemental Table 1**. Spot blotch DSR of tested global wheat germplasms.

**Supplemental Table 2**. List of leaf spot-resistant wheat lines identified in this study.

**Supplemental Table 3**. SNPs significantly associated with spot blotch resistance.

**Supplemental Table 4**. Information on annotated genes within the *Qsb.hebau-1BL*

interval in the Chinese Spring reference genome.

**Supplemental Table 5**. Haplotype analysis on 9 SNPs within the *Qsb.hebau-1BL* physical interval.

**Supplemental Table 6**. dCAPS markers designed based on SNPs associated with *Qsb.hebau-1BL*.

**Supplemental Table 7**. Sequencing information for RNA samples in this study.

**Supplemental Table 8**. Expression profile of pathogenesis-related protein (*PR*) genes in wheat resistance to spot blotch.

**Supplemental Table 9**. Information on DEGs in the comparison of “R522 (*Qsb.hebau-1BL*)_BS vs R522_CK” annotated in the KEGG pathway of “MAPK signaling pathway -plant”.

**Supplemental Table 10**. Expressions of candidate genes within the *Qsb.hebau-1BL* interval in wheat resistance to spot blotch.

**Supplemental Table 11**. Variations of the *TaNADPO* gene among wheat lines with or without *Qsb.hebau-1BL*.

**Supplemental Table 12**. dCAPS markers designed based on SNPs in the promoter region of the *TaNADPO* gene.

**Supplemental Table 13**. Primers used for qRT-PCR assay in this study.

**Supplemental Table 14**. Primers used for amplification of the *TaNADPO* gene.

