## Supplemental Figures for "Wheat *TaNADPO* promotes spot blotch resistance"

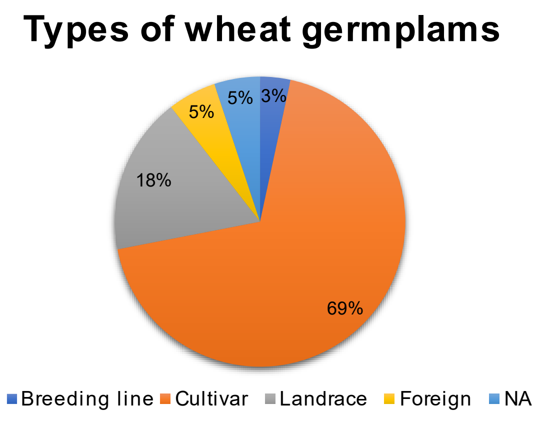


**Supplemental Figure 1.** Types of wheat germplasms in the studied natural population.

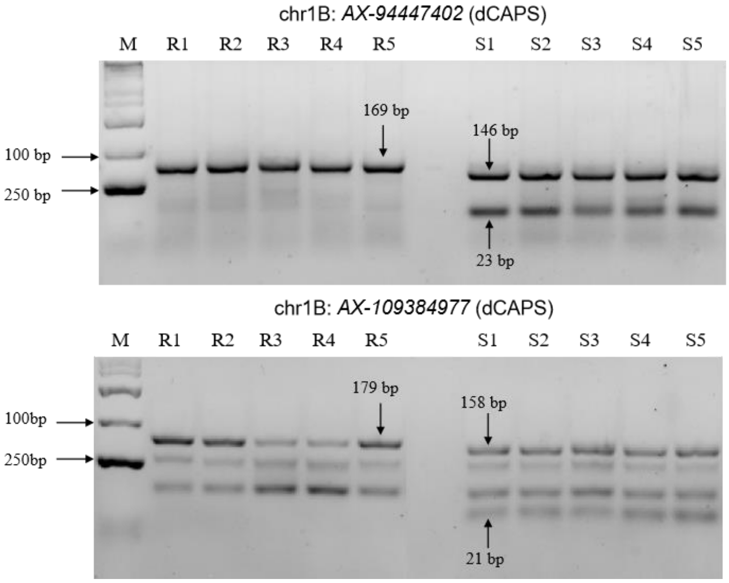


**Supplemental Figure 2.** Validation of dCAPS markers designed based on SNPs within the *Qsb.hebau-1BL* locus.

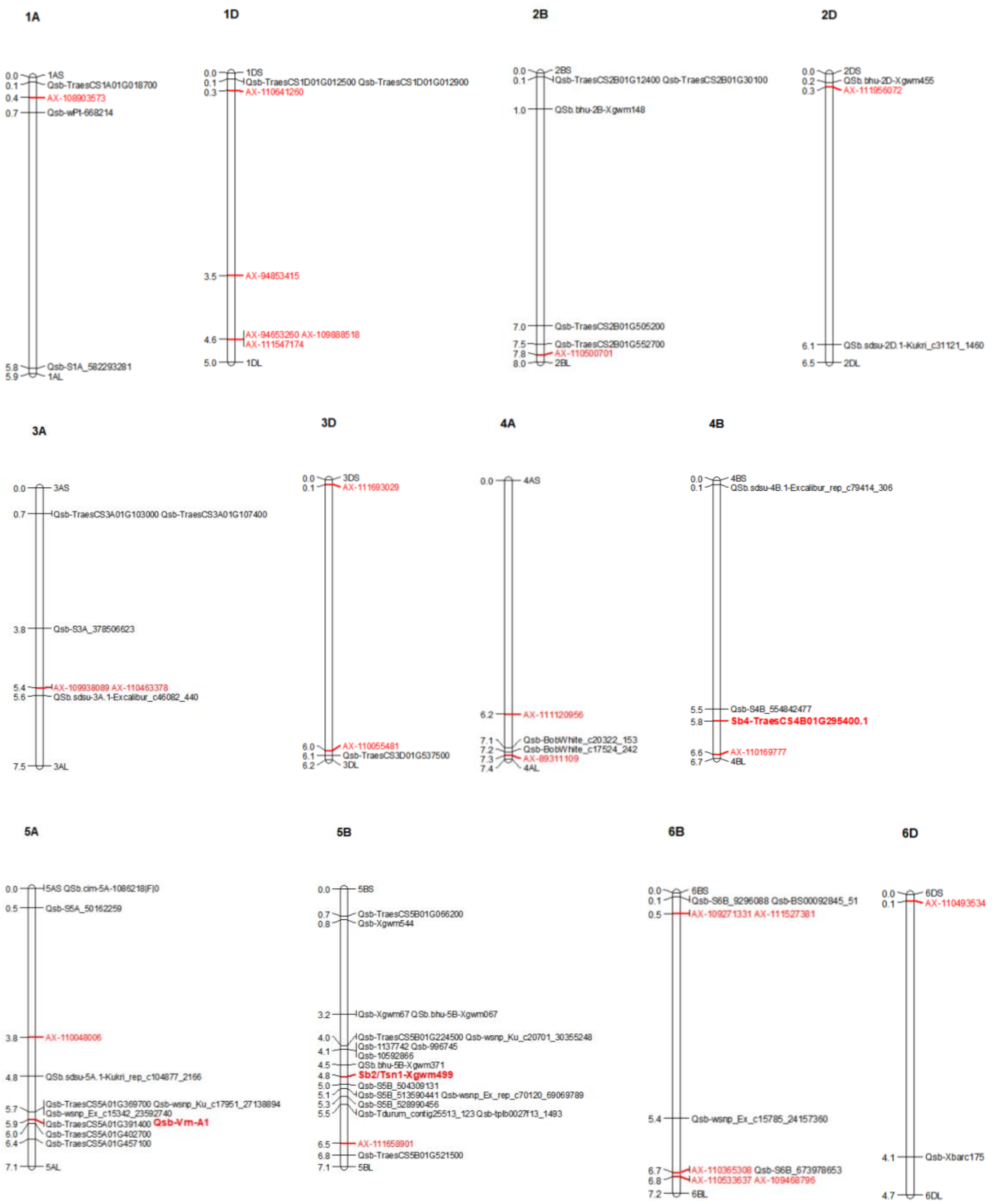


**Supplemental Figure 3.** Distribution map displaying associated SNPs and previously reported spot blotch resistant QTLs on chromosomes other than Chr1B.

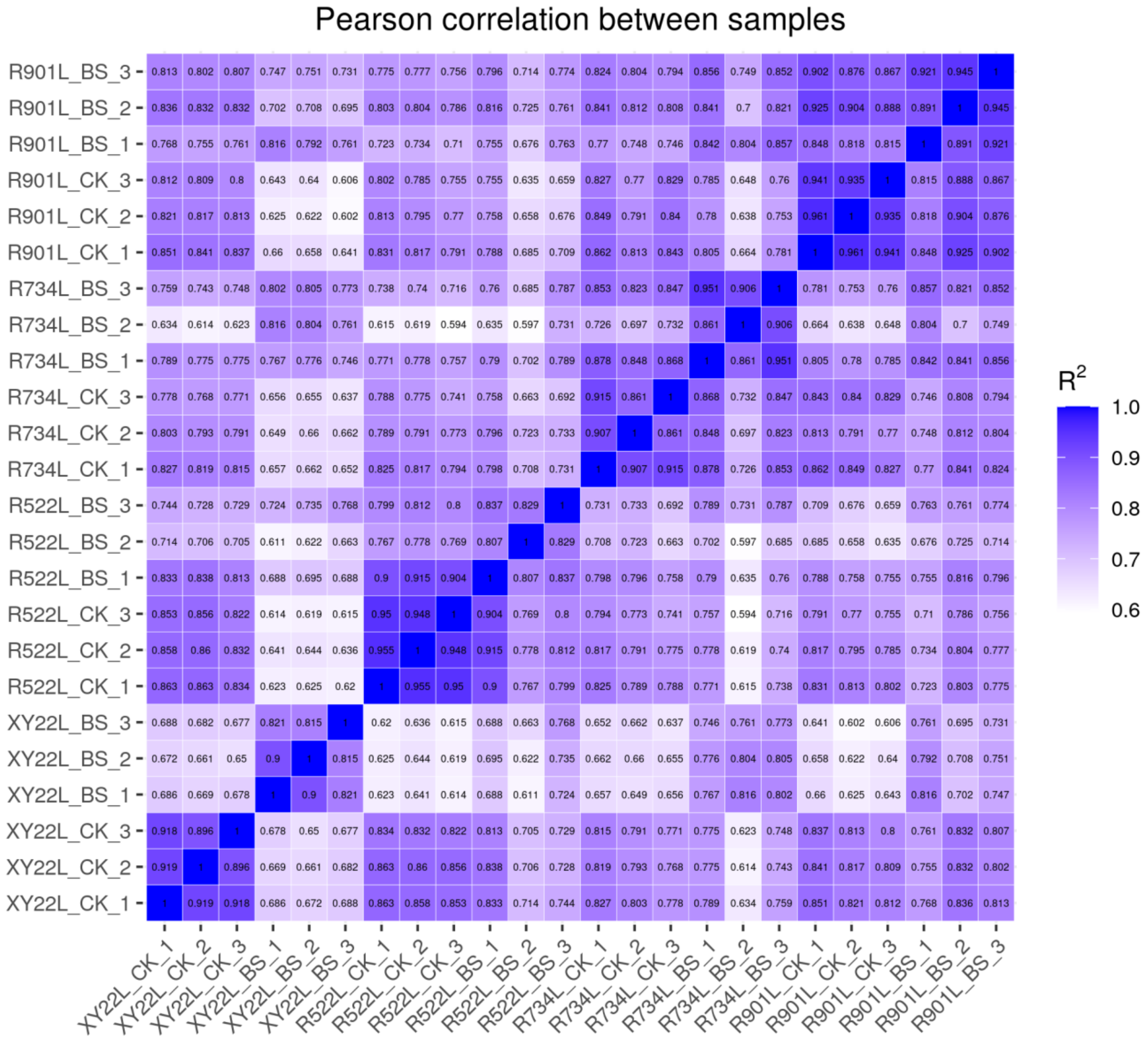


**Supplemental Figure 4.** Principal component analysis (PCA) of the overall transcript expressions for each sample in the transcriptome.

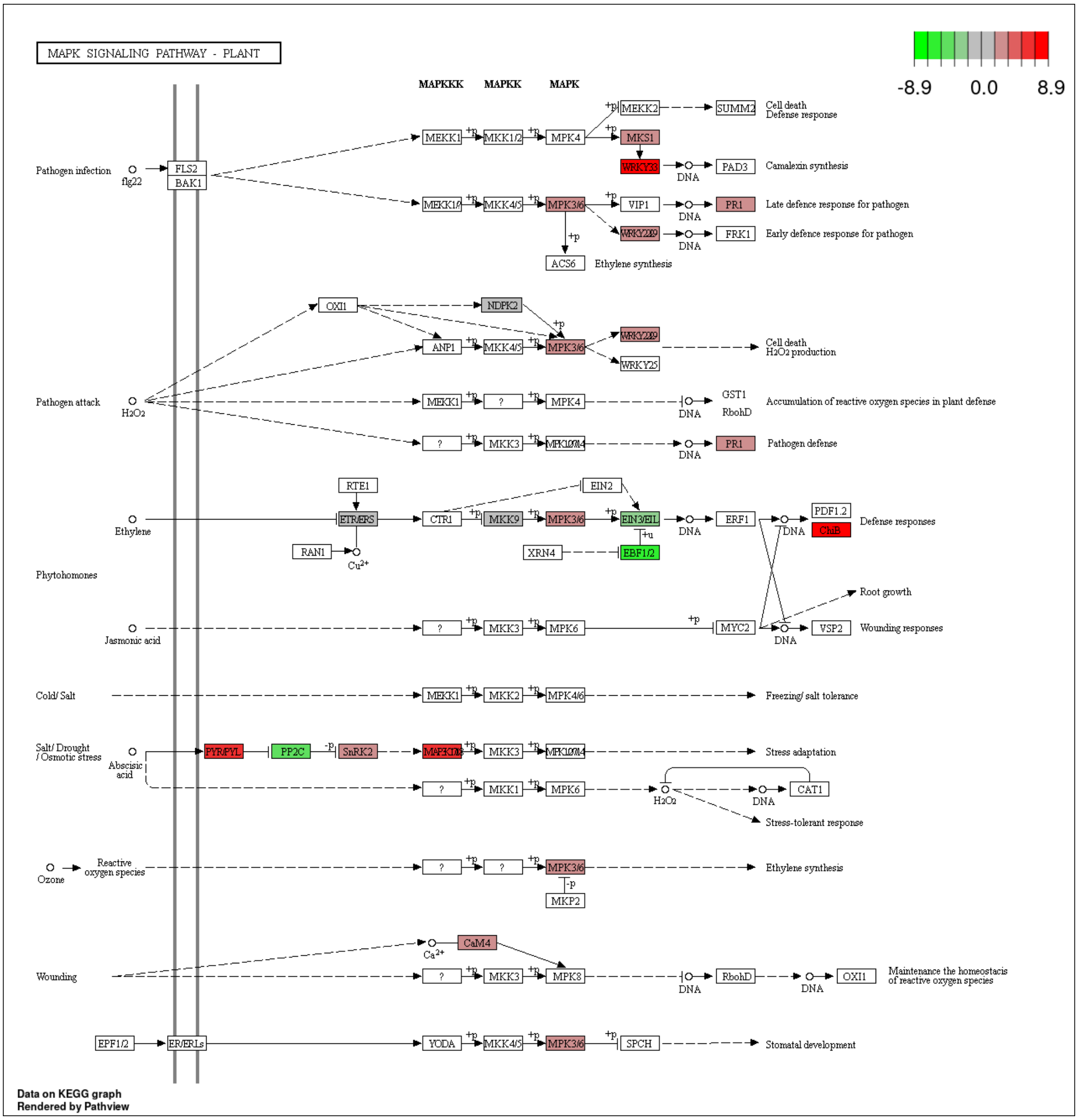


**Supplemental Figure 5.** DEGs in comparison of “R522L_BS vs R522L_CK” enriched in the KEGG pathway of “MAPK signaling pathway – plant”.

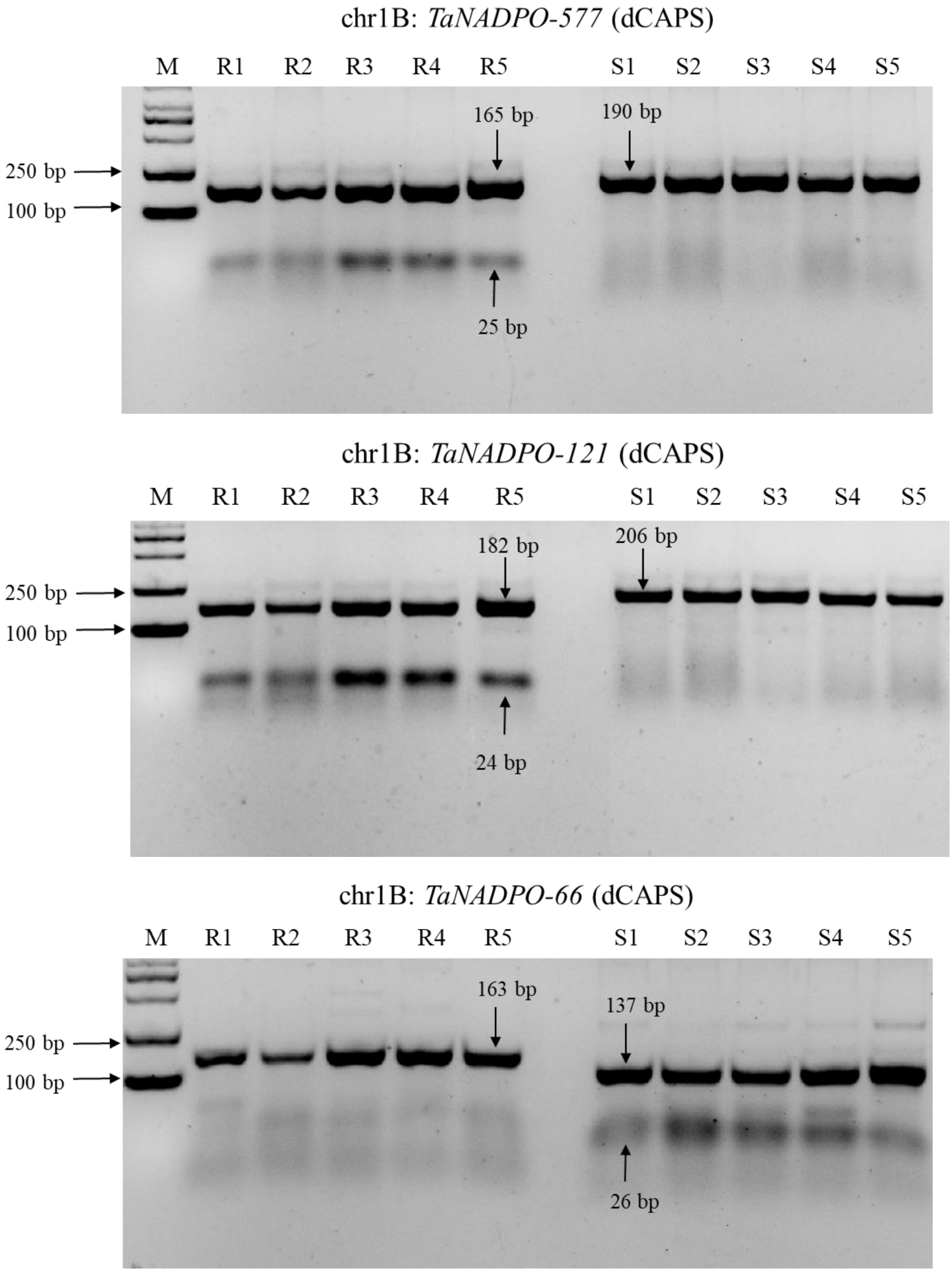


**Supplemental Figure S6.** Development and validation of three dCAPS markers based on variations in the *TaNADPO* gene.

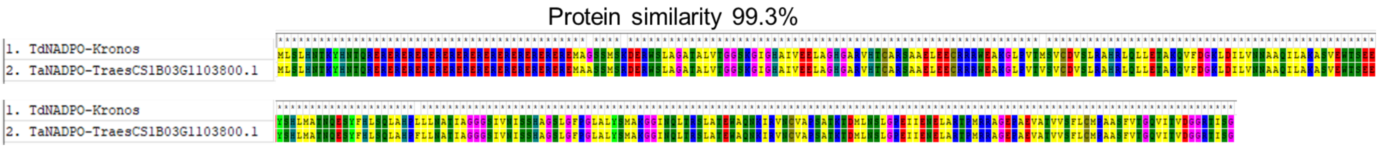


**Supplemental Figure S7.** Protein sequence alignment of TaNADPO with TdNADPO.

A B

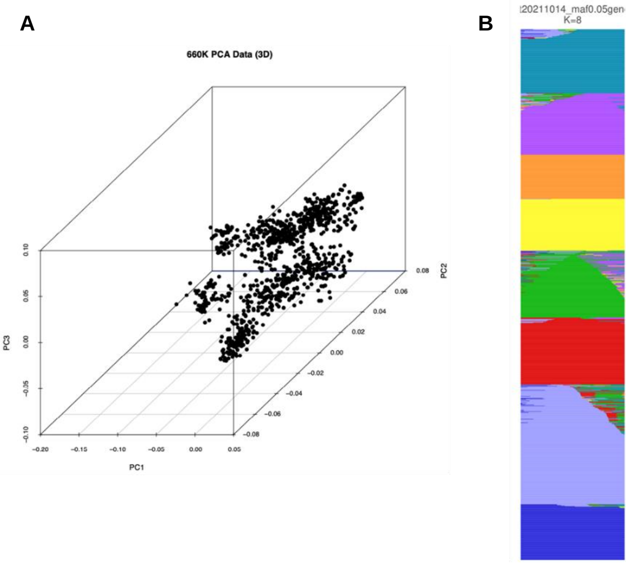


**Supplemental Figure S8.** Population structure analysis of the studied natural population.
**(A)** Principal Component Analysis (PCA) diagram.
**(B)** Population structure diagram.

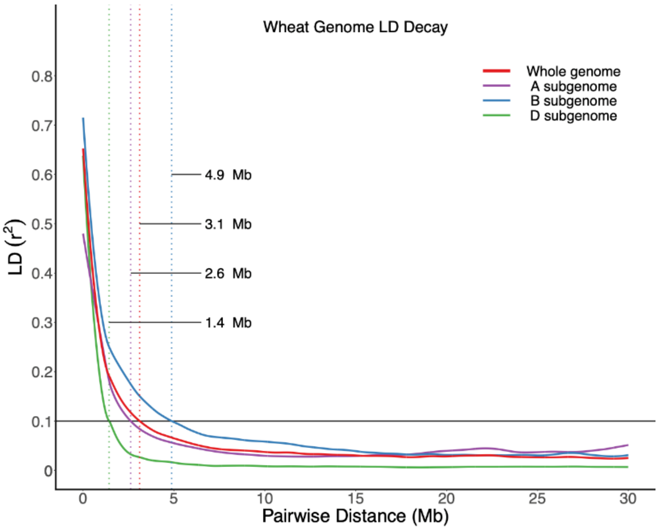


**Supplemental Figure 9.** Linkage disequilibrium (LD) decay map of the SNPs utilized in this study.
